# Early reduction and impaired targeting of myelin-associated glycoprotein to myelin membranes in Huntington’s disease

**DOI:** 10.1101/2025.10.15.682629

**Authors:** Adel Boudi, Ellen Sapp, Yvonnie Y Li, Kai Lee Shing, Kimberly Kegel-Gleason, Tiziana Petrozziello, Ghazaleh Sadri-Vakili, Neil Aronin, Marian DiFiglia, Xueyi Li

**Affiliations:** Department of Neurology, Massachusetts General Hospital, 114 the 16 street, Charlestown MA 02129, USA; RNA Therapeutics Institute, University of Massachusetts Chan Medical School, 368 Plantation Street, Worcester MA 01605, USA

**Keywords:** Huntington’s disease, white matter, oligodendrocytes, endosomes, myelin-associated glycoprotein

## Abstract

**Background:** Huntington’s disease (HD) is a hereditary life-threatening disease marked by progressive neuronal loss and atrophy of grey matter structures, particularly the caudate putamen. Brain imaging studies have revealed that the degradation of the white matter occurs many years prior to symptomatic onset and neuronal loss, suggesting that the decay of brain white matter is an active contributor to the disease progression. However, the mechanisms by which the HD mutation triggers white matter loss is not well understood.

**Methods:** Western blot, immunohistochemistry, and electron microscopy were conducted to assess white matter pathology and explore the relevant mechanisms in CAG140 knock-in mice, which express the HD protein in the same way as patients suffering from HD and thus biologically replicate HD in human.

**Results:** Western blot analysis of proteins localized at different layers of the myelin coat revealed that the myelin-associated glycoprotein (MAG), which is localized at the innermost layer of the myelin coat and essential for maintaining the periaxonal space and the integrity of the myelin sheath, manifested as an early and progressive decline in HD mouse caudate putamen. The loss of MAG was detected at myelinated axons and in fiber bundles in HD mouse brains at an age when the abundance of myelinated axons was normal. Fluorescence immunohistochemical studies found that MAG labeling was concentrated in the soma of a subset of oligodendrocytes, which expressed breast carcinoma amplified sequence 1, a marker for new oligodendrocytes. While their abundance was normal, new oligodendrocytes in HD mouse caudate appeared to be impeded in acquiring the expression of MAG and in targeting MAG away from perinuclear punctate structures to processes, signs of impaired maturation. Compared with those in wildtype mouse brains, oligodendrocytes in HD mouse brains had a reduced abundance of small vesicles whereas an increased abundance of large punctate structures in the perinuclear region, implying defective generation of small vesicles transporting MAG from large punctate structures in the soma to processes. The MAG-containing perinuclear punctate structures were negative for proteins specifying trans-Golgi networks, early endosomes, or exosomes but had a minor portion labeled with lysosome-associated membrane protein 1, indicating that the structures where MAG accumulates in the soma are derived from the late endosomal lysosomal compartment.

**Conclusions:** Our study suggests that the decay of the brain white matter in Huntington’s disease involves a deficit in trafficking of myelin-associated glycoprotein, preventing its proper delivery from the soma of oligodendrocytes to myelin-forming processes.

## Introduction

Huntington’s disease (HD) is associated with progressive degeneration of neurons and atrophy of grey matter structures particularly the caudate putamen. Brain imaging studies have revealed that the white matter, which comprises myelinated axons and glial cells, undergoes deterioration in the brain of HD patients at early and presymptomatic stages (*1*). Deficits in the myelin formation have been identified in postmortem human HD brains (*2, 3*), supporting the hypothesis of a “demyelination” process that results in severe loss of brain white matter in HD (*4*). In HD mouse models (YAC128, BACHD, and CAG knock-in) expressing the human HD transgene or mouse huntingtin (*Htt*) with a pathogenic CAG expansion, evidence indicates that white matter degeneration precedes neuronal degeneration (*5-9*). Collectively, these lines of evidence suggest that the degradation of the brain white matter is unlikely a secondary effect of neuronal degeneration but an active contributor to disease progression in HD. However, the mechanisms by which the HD mutation drives the white matter degeneration remain poorly defined.

In the brain, myelin is formed by the flat sheet-like processes of oligodendrocytes, which arise from oligodendrocyte precursor cells (OPCs) through a well-defined differentiation program (*10, 11*). Throughout life, new oligodendrocytes are generated to maintain existing myelin, repair damage, and myelinate previously unmyelinated axons in response to experiences (*10-12*). Single-cell RNA sequencing analysis of postmortem human and mouse HD brains has revealed that the mutant huntingtin (mHtt) impairs oligodendrocyte maturation and alters lipid and glucose metabolism in oligodendrocytes (*13*). These changes may underscore the re-myelination defects observed in adult HD brains. Although it is an intrinsic property of oligodendrocytes, myelination requires neuronal cues to promote the delivery of myelin building blocks to the myelin membranes (*14*). On the other hand, oligodendrocytes release exosomes that influence neuronal firing rate, signal transduction, and gene expression (*15*). These findings highlight the importance of the neuron-oligodendrocyte communication in myelination. However, little is known about whether and how the HD mutation interferes with neuron-oligodendroglia interactions.

Myelin-associated glycoprotein (MAG), a transmembrane glycoprotein present in both central and peripheral nervous systems, is a key player in crosstalk between oligodendrocytes and neurons. Localized at the innermost layer of the myelin sheath, MAG interacts with gangliosides and Nogo-receptors on the opposing axolemma (*16-18*). While it itself is not required for the initial myelination, MAG is essential for maintaining the periaxonal space between axons and the innermost layer of myelin membranes (*19, 20*). Loss of MAG reduces axon diameter and causes axonal atrophy (*21*). Mutations of the MAG gene are linked to schizophrenia, autosomal recessive spastic paraplegia type 75, and Pelizaeus-Merzbacher disease-like disorder (*22-25*). Additionally, an early decline in the level of MAG protein has been found in various neurological disorders, including multiple sclerosis and Alzheimer’s disease (*26-29*).

Myelin-associated proteins are sequentially expressed during the maturation of oligodendrocytes; MAG is expressed earlier than myelin basic protein (MBP), myelin proteolipid protein (PLP) and myelin oligodendrocyte glycoprotein (MOG), which are localized at compact and outermost myelin membranes, respectively (*30, 31*). Oligodendrocyte maturation involves the remodeling of the plasma membrane into distinct domains, namely the soma, processes, and myelin membrane sheets, and coordinated exocytosis and endocytosis to direct myelin components to membrane sheets (*10, 11, 32*). This sorting relies on vesicle machineries employed for generating apical and basolateral membrane domains in polarized epithelial cells (*33, 34*), among which Rab11 plays an important role (*35*). mHtt has been shown to disturb vesicle trafficking in multiple pathways and dampen Rab11 function (*36, 37*). Taken together, these observations suggest that the HD mutation may interfere with the targeting of MAG as well as other myelin components to axon-ensheathing membranes, thereby contributing to white matter pathology in HD.

In this study, we investigated white matter pathology in HDQ140 knock-in mice. Among the myelin-associated proteins examined, MAG exhibited an early and progressive reduction in its level in the caudate putamen. Localization of MAG was diminished in processes ensheathing axons and accumulated in perinuclear structures within a subset of oligodendrocytes expressing breast carcinoma amplified sequence 1 (BCAS1), a marker of newly generated oligodendrocytes. Imaging analysis further revealed that these MAG-positive structures were significantly enlarged in HD mouse brains compared to those in WT mice. This subcellular distribution pattern of MAG did not affect the abundance of myelinated axons. Taken together, our study suggests that white matter defects in HD involves impaired targeting of MAG from the oligodendrocyte soma to myelin membranes.

## Materials and Methods

### Animals

Animals were housed in male and female groups and maintained in a pathogen-free environment under regular 12-hour light/12-hour dark cycle with food and water *ad libitum*. All mice were genotyped by PCR as described. Experimental procedures were carried out following the NIH Guide for the Care and Use of Laboratory Animals and approved by the Institutional Animal Care and Use Committee of Massachusetts General Brigham (#2004N000248). Although there are sex differences in the severity of symptoms, both males and females are attacked by HD (*38*). Therefore, mice of both sexes were used for studies.

### SDS-PAGE and Western blot analysis

For studies of mouse brain, small pieces of striatum and cortex were dissected from fresh WT, HDQ140/Q140, and zQ175 mouse brains. For studies of human postmortem brain, frozen blocks from prefrontal cortex and caudate putamen were dissected. Bain pieces were homogenized through a dounce homogenizer on ice in pre-cooled homogenization buffers containing 10mM HEPES (pH7.2), 1 mM EDTA, 250 mM sucrose, 1mM NaF, 1mM Na3VO4 and a cocktail of protease inhibitors (Roche). Crude homogenates were sonicated for 10 seconds, and the concentration of proteins in each sample was determined using the Bradford method (Bio-Rad). An equal amount (10 μg) of proteins from each sample was separated by SDS-PAGE and transferred onto nitrocellulose membranes with a Transblot Turbo apparatus (Bio-Rad). Blots were cut horizontally to increase the number of antibodies analyzed per gel, blocked in 5% non-fat milk dissolved in Tris buffered saline containing 0.1% Tween-20 (TBST), and then incubated with primary antibodies overnight at 4°C with gentle agitation. After 3 washes in TBST, blots were incubated with peroxidase-conjugated secondary antibodies (Jackson ImmunoReasearch). After incubation with secondary antibodies, blots were washed 3 times in PBST, and protein bands were visualized with SuperSignal WestPico Plus Chemiluminescent kit (Pierce) and ChemiDoc XRS imager (Bio-Rad). Densitometry was performed using the NIH ImageJ/Fiji by manually circling the protein bands of interest to measure area and average intensity, which were used for calculating total signal intensity by multiplying them. Primary and secondary antibodies, working dilutions, and suppliers used for Western blot analysis were summarized in **Table 1**. Primary and secondary antibodies were diluted in TBST containing 5% non-fat milk.

**Table 1.**
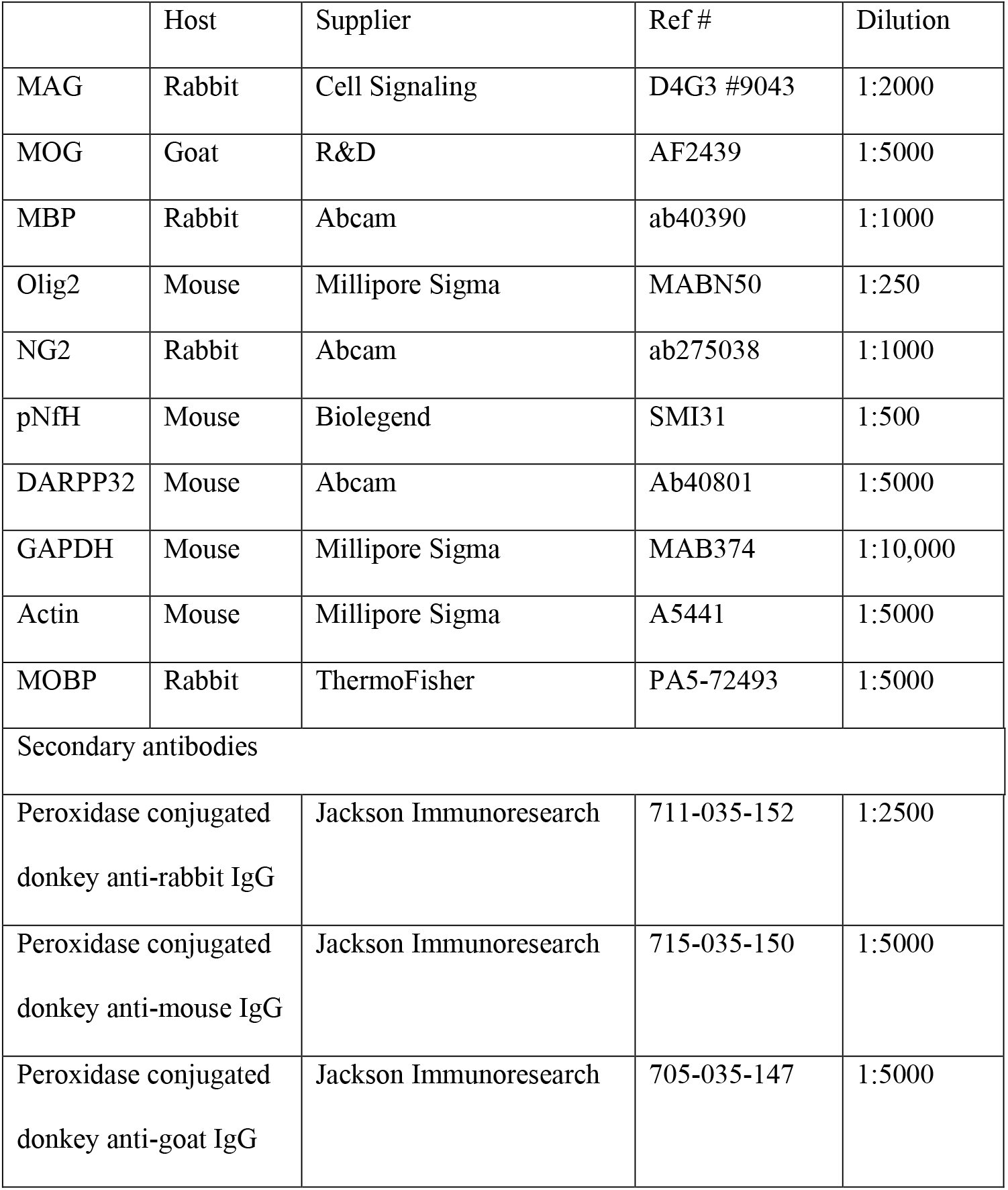
Primary and secondary antibodies used for Western blot.

### Immunofluorescence microscopy and image analysis

Mice at age 4 to 5 months were deeply anesthetized and transcardially perfused with 50 ml of physiological phosphate buffer solution (PBS) followed by 50 ml of 3.7% paraformaldehyde (Sigma-Aldrich) in PBS. Fixed brains of WT and HD (HDQ140/Q140 and zQ175) mice were cut into 30-μm-thick coronal sections on a cryostat (Leica CM1950). A series of 3 consecutive brain sections from each of 3 WT and 3 HD mice were used for immunolabeling, which was performed with free-floating brain sections. Sections were washed in PBS at room temperature and then permeabilized in PBS containing 5% normal donkey serum, 0.2% Triton X-100, and 1% bovine serum albumin for 1 hour and then incubated with primary antibodies at 4°C with gentle agitation for 48 hours. After washes in PBS containing 0.05% Triton X-100, brain sections were incubated with fluorescent dye-conjugated secondary antibodies (1:500, Jackson ImmunoResearch) at room temperature for 2 hours. Cells in brain sections were identified by staining nuclei with Hoechst 34580 (Invitrogen). Brain sections were mounted onto glass slides with ProLong Gold (Invitrogen) for microscopy. Primary and secondary antibodies, working dilutions, and suppliers were summarized in **Table 2**. All antibodies used for immunofluorescence labeling were diluted in PBST containing 5% normal donkey serum, 0.05% Triton X-100, and 1% bovine serum albumin.

**Table 2.** Primary and secondary antibodies used immunofluorescence microcopy.

|  | Host | Supplier | Ref # | Dilution |
| --- | --- | --- | --- | --- |
| MAG | Rabbit | Cell Signaling | D4G3 #9043 | 1:2000 |
| pNfH | Mouse | Biolegend | SMI31 | 1:500 |
| BCAS1 | Rabbit | Abcam | ab275038 | 1:1000 |
| MOBP | Rabbit | ThermoFisher | PA5-72493 | 1:5000 |
| Golgin97 | Mouse | Santa-Cruz | sc-59820 | 1:100 |
| EEA1 | Mouse | BD Biosciences | 610456 | 1:200 |
| LAMP1 | Rat | Invitrogen | 14-1071-82 | 1:200 |
| CD63 | Mouse | Abcam | Ab8219 | 1:500 |
| Secondary antibodies |  |  |  |  |
| C3 or Rhodamine conjugated donkey anti-rabbit IgG |  | Jackson ImmunoResearch | 711-035-152 | 1:000 |
| Alexa488 or Bodipy conjugated donkey anti-mouse IgG |  | Jackson ImmunoResearch | 715-035-150 | 1:500 |

Digital images were acquired through an LSM 880 confocal microscope (Carl Zeiss) using 405, 488, 561, and 633 nm lasers. Images used for quantitative analysis of the expression levels of proteins and oligodendrocyte mature stages were captured with the same settings, including laser strength, pinhole, signal gain, and offset. Quantitative analyses of confocal images were conducted with the NIH ImageJ/Fiji. Researchers who collected images and conduced image analyses were blinded to genotypes. For measuring signal intensities of immunoreactivities for MAG and pNfH, respectively, the soma of each of MAG-labeled oligodendrocytes in the caudate putamen area in each imaged brain section was manually circled, and the area and average signal intensity were measured and used for calculating total signal intensity of MAG immunoreactivities in the soma. The same contour surrounding the soma of the oligodendrocyte was placed in the center of a fiber bundle in proximity to the soma, and the average signal intensity of MAG as well as pNfH in the fiber bundle was measured. The ratio of total signal intensity of MAG in the soma to that in the nearby fiber bundle was calculated and used as a criterion for determining the accumulation of MAG in the soma. The expression level of pNfH was expressed as the signal intensity of pNfH in the fiber bundle. For quantitative analysis of the maturation stages of new oligodendrocytes in WT and HD mouse caudate putamen, cells labeled with MAG only or BCAS1 only or with both MAG and BCAS1 in the cell body in each imaged brain section was counted. Based on the signal strength of MAG immunoreactivities relative to BCAS1 immunoreactivities, cells containing both MAG and BCAS1 in the soma, could be further classified into 3 subgroups: low MAG levels whereas high BCAS1 levels, similar MAG and BCAS1 levels, and high MAG levels whereas low BCAS1 levels. The percentage of each of the 5 populations of oligodendrocytes in each of WT and HD mice was calculated. For analyzing the size of MAG-labeled structures in the soma of oligodendrocytes, images were collected with the same magnification and opened. After appropriately adjusting the signal threshold of images opened with the NIH ImageJ/Fiji, the “Analyze Particles” tool was applied to measure MAG-labeled structures in the soma. The “abnormally large particles” identified with the NIH ImageJ/Fiji were further examined manually to ascertain that they were not derived from fusion of two or more small “particles”. The value of the measured cross-sectional area of each “particle” was converted to μm^2^ according to the original resolution of the image.

### Immuno-gold electron microscopy and quantitative analysis

Slices (approximately 3mm x 2mm x 1mm) were cut from dorsal caudate underlying primary motor cortex of each mouse brain (n=3 mice per genotype) and further cut into small blocks. Tissue blocks from each mouse were cryo-protected by an overnight infiltration with 2.3 M sucrose in PBS at 4 °C and then mounted on aluminum pins and frozen in liquid nitrogen. Cryosections of 60 – 80 nm were cut on a Leica EM FCS at –80 °C, collected on formvar-coated nickel grids, and floated in PBS with 0.02% sodium azide as a preservative. Immunogold labeling with the protein A-gold method was done as standard procedures. In brief, grids were blocked with 1% BSA in PBS and incubated with anti-MAG antibody for 1 hr at room temperature followed by incubation with protein-A conjugated with 15 nm gold particles for 1 hr. After several rinses on drops of distilled water, the grids were floated on drops of Tylose® cellulose and uranyl acetate for 10 min on ice, collected on loops and allowed to dry. The grids were examined at 80 kV in a JEOL JEM 1011 transmission electron microscope. Images were taken of areas with clusters of cross-sectioned myelinated axons using an AMT digital imaging system (Advanced Microscopy Techniques, Danvers, MA). The number of myelinated axons with or without immunogold labeling as well as the number of gold particles at each myelinated axon was quantified by observers blinded to genotypes in 10 – 11 images per mouse. The mean number of gold particles per axon was determined for each group.

### Statistics

Data were expressed as Mean±SD. Statistical significance was determined by two-tailed unpaired Student’s t-test. A P value less than 0.05 was considered significant.

## Results

### Mutant huntingtin is expressed and forms aggregates in brain oligodendrocytes

Prior studies have reported increased oligodendrocyte proliferation by impaired maturation in postmortem human and mouse HD brains (*6, 13*). The finding that genetic abrogation of mHtt specifically in oligodendroglia attenuates white matter pathology in HD supports an intrinsic contribution (*39*). To further address this, we sought to confirm the expression of mHtt proteins in oligodendroglia. Brain sections from HDQ140/Q140 knock-in mice, which express mHtt under the endogenous mouse *Htt* promoter, were immunolabeled with the 2B7 antibody recognizing the first 17 amino acids of Htt. Oligodendrocytes were identified by co-labeling with MAG. Fluorescent microscopic analysis revealed that Htt immunoreactive signals were hardly detectable in the soma but visible in processes containing MAG (**Figure 1A**). To corroborate these findings, sections were co-labeled with MAG and Htt MAB2166 antibodies, which recognizes an epitope within amino acid 181-810 of Htt. Despite being relatively low when compared with those in nearby cells, MAB2166-labeled signals were clearly present in the soma of MAG-positive oligodendrocytes (**Figure 1B**). We next examined whether mHtt forms aggregates in oligodendrocytes. Brain sections of zQ175 mice, line derived from HDQ140 knock-in mice and developing disease-relevant phenotypes with only one allele mutated (*40, 41*), were labeled with the S830 antibody, which detects mHtt aggregates in the striatum of zQ175 mice as early as age 6 weeks (*42*). Consistent with prior findings (*42*), nuclear mHtt aggregates were widely distributed throughout the caudate putamen of 12-month-old zQ175 mice (**Figure 1C**). Small mHtt aggregates were detected in the cytoplasm of MAG-positive oligodendroglia (**Figure 1C**). Collectively, these findings indicate that mutant Htt is bona fide expressed in oligodendrocytes and forms aggregates.

**Figure 1.**
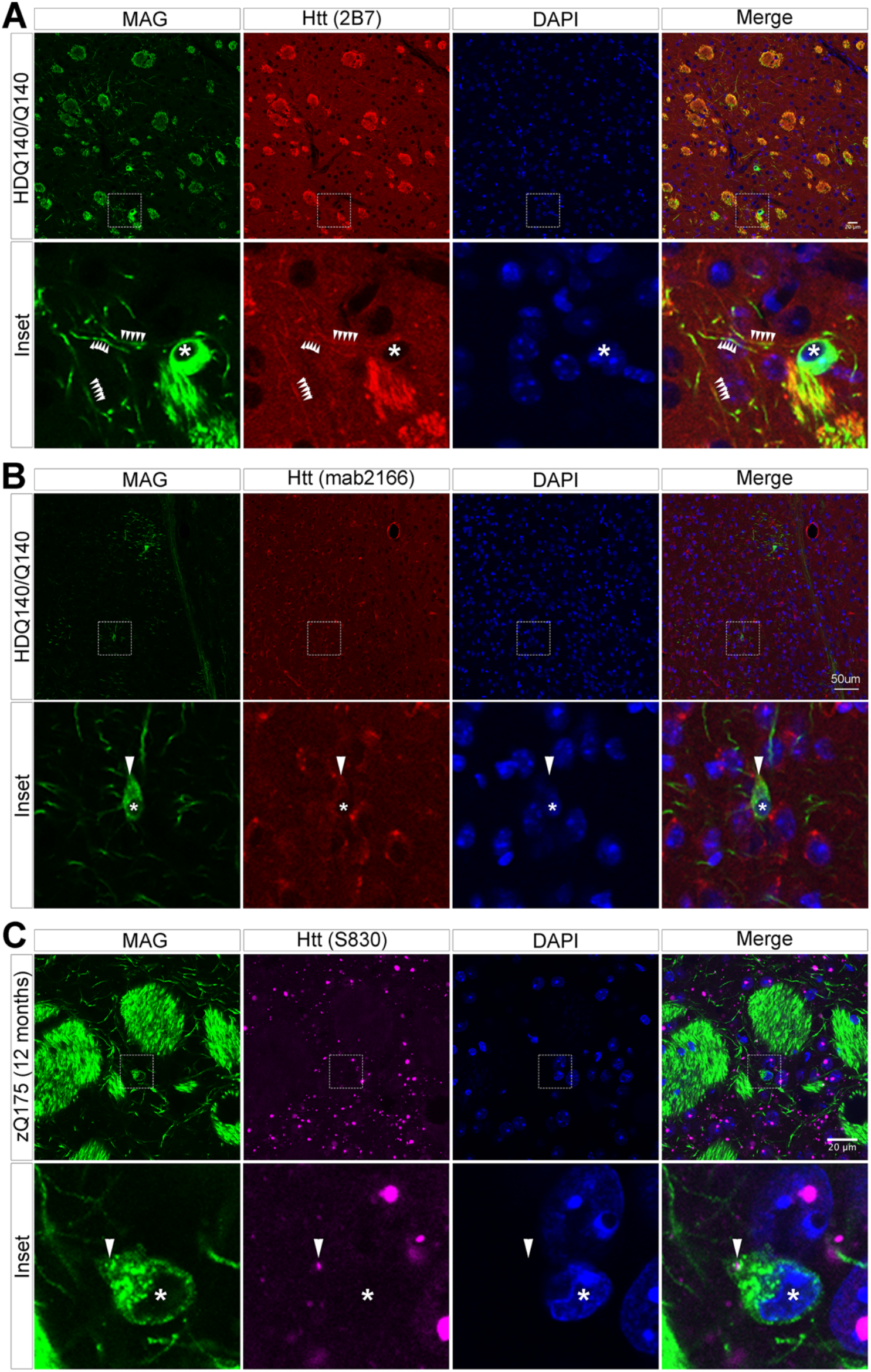
Mutant huntingtin is expressed and forms aggregates in oligodendrocytes. A series of 3 consecutive coronal sections cutting through the caudate putamen of 4 to 5 months old HDQ140/Q140 mice (**A** and **B**) or 6 months old zQ175 mice (**C**) were processed for labeling with antibodies for MAG combined with antibodies for Htt 2B7 (A), or MAB2166 (B), or S830 (C). After incubation with fluorescent dye-conjugated secondary antibodies, brain sections were mounted onto glass slides for microscopy. Cells in brain sections were identified by staining nuclei with DAPI. Shown are confocal images. Boxed regions in each image in (**A, B** and **C**) were enlarged and shown as Inset below the corresponding image. Arrowheads in (**A**) point to segments of oligodendroglia processes, in (**B**) indicate the cell body of an oligodendrocyte, and in (**C**) point to the position of a cytoplasmic Htt aggregate in an oligodendrocyte. The stars in (**A, B** and **C**) identify the nuclear region of an oligodendrocyte.

### Altered distribution of MAG in oligodendroglia in HD mouse brains

During our analyses of mHtt expression in oligodendrocytes, we noticed a marked reduction of MAG immunoreactivities in fiber bundles in the caudate putamen of HDQ140/Q140 mice (**Figure 1**). To further characterize this phenotype, we compared MAG distribution pattern in the caudate putamen of WT and HDQ140/Q140 mice at age 4 to 5 months. Brain sections of WT and HDQ140/Q140 mice were labeled with antibodies against MAG along with antibodies against phosphorylated neurofilament heavy chain (pNfH) to detect MAG in the soma of oligodendrocytes and in myelin membranes (**Figure 2A**). We quantified the distribution of MAG by calculating the ratio of MAG signal intensity in oligodendrocyte soma relative to adjacent fiber bundles. While its average was <1 in WT mice this ratio was >3 in HdhQ140/Q140 mice (**Figure 2B**), suggesting that the delivery of MAG from compartments in the soma to myelin membranes is impaired in HD mouse brains. We then determined whether the reduction of MAG in fiber bundles was a consequence of axonal damage by analyzing pNfH immunoreactive signals in the fiber bundles. Our results showed no genotype-dependent difference, indicating preserved axonal integrity (**Figure 2A, 2C**). Together, these data suggest that the reduction of MAG in fiber bundles in HDQ140/Q140 mice reflects defective targeting from oligodendrocyte soma to myelin membranes rather than secondary loss due to axonal damage.

**Figure 2.**
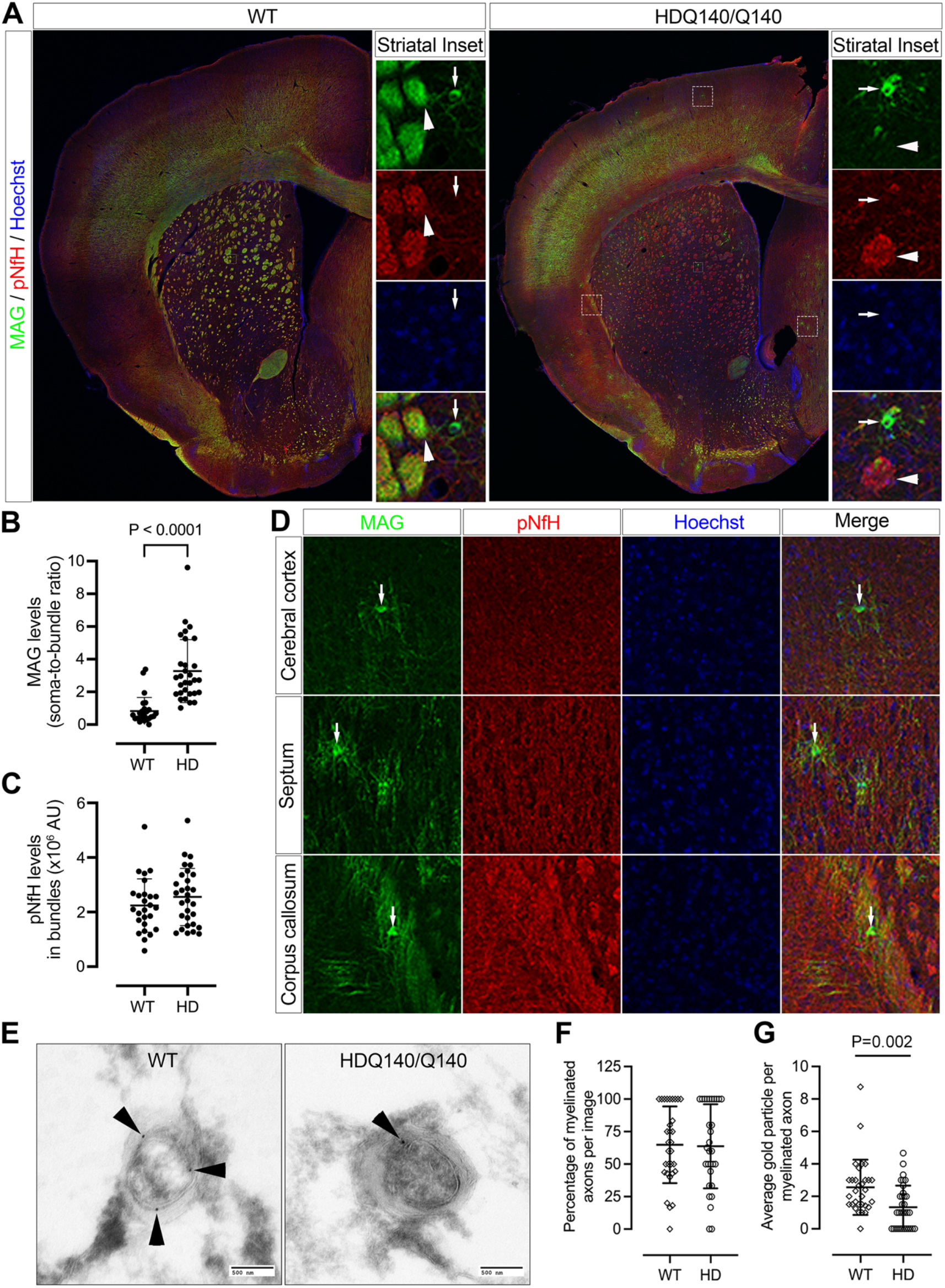
MAG is diminished at myelinated axons and accumulates in the cell body of a subset of oligodendrocytes. **A**) Immunofluorescent labeling of WT and HDQ140/Q140 mice brains sections with antibodies for MAG combined with antibodies for pNfH to detect MAG in the soma of oligodendrocytes and in a closely nearby pNfH-labeled fiber bundle, respectively. Shown are confocal merged images of one-half brain section of a WT mouse and a HD mouse. Cells in brain sections were identified by staining nuclei with DAPI. Boxed regions in striatal areas were enlarged and shown in separate channels to the right of the corresponding merged image. The caudate putamen region in each imaged half brain section was chosen for densitometrical analyses of MAG signals in the soma and in a nearby fiber bundle (**B**) and pNfH signals in the corresponding fiber bundle (**C**). Each symbol represents an oligodendrocyte having intense MAG signals in the soma (**B**) and one fiber bundle (**C**). **D**) Shown are images of the enlarged boxed regions in the cerebral cortex, septum and/or corpus callosum areas in imaged brain sections (**A**). Arrowheads in Striatal Insets (**A**) point to a fiber bundle close to an oligodendrocyte containing intense signals of MAG in the soma, whereas arrows in Striatal Insets (**A**) and in (**D**) point to oligodendrocytes containing intense MAG signals in the soma. Electron microscopic analysis of MAG-labeled ultrathin sections of mouse caudate tissues (**E**) followed by quantitative analysis of myelinated axons (**F**) and gold particles at myelinated axons (**G**). Arrowheads in photographs (**E**) indicate MAG labeled with gold particles at myelinated axons. Statistical significance was determined by unpaired two-tailed Student’s t-test. Data are mean±SD (n=3 mice per genotype, 3 sections/mouse, age of animal: 4 to 5 months).

Oligodendrocytes with intense MAG signals in the soma were not restricted to the caudate putamen but were also observed in other brain regions, including the cerebral cortex, the medial septal nucleus, and the corpus callosum (**Figure 2D)**. To determine whether this abnormal distribution was displayed by other myelin-associated proteins, brain sections from WT and HDQ140/Q140 were labeled with antibodies for myelin-associated oligodendrocyte basic protein (MOBP). Like MAG, MOBP immunoreactivity was markedly decreased in fiber bundles in the caudate putamen of HDQ140/Q140 mice (**Figure S1**). However, unlike MAG, MOBP did not accumulate in perinuclear structures within oligodendrocytes. (**Figure S1**). Taken together, these data suggest that oligodendrocytes in HD mouse brains have a deficit in targeting MAG from compartments in the cell body to myelin-forming processes.

### MAG is diminished at myelinated axons in HDQ140/Q140 mice

MAG localizes at periaxonal myelin membranes, where it plays an essential role in the maintenance of the space between the innermost layer of myelin membranes and the axonal surface (*19, 20*). To corroborate that the loss of MAG occurred at processes that wrap around axons, we conducted electron microscopic studies to corroborate that. Dorsal caudate tissue blocks of WT and HDQ140/Q140 mice were cut into ultrathin sections for immunogold labeling to detect MAG. In cross-sectioned fibers, gold particles labeling MAG appeared at the myelin membranes adjacent to the axolemma (**Figure 2E**). Quantitative analysis of randomly selected fields revealed that the abundance of myelinated axons was not different between WT and HDQ140/Q140 mice. However, the average number of gold particles per myelinated axon was significantly reduced in HDQ140/Q140 mice compared to WT mice (**Figure 2F, 2G**), suggesting that the reduction of MAG occurs prior to overt myelin breakdown. These data support the above idea that the targeting of MAG to myelin-forming processes is impaired in HD.

### Reduction of MAG in HD mice is an early and progressive phenotype

Having shown the reduction of MAG in HD mouse brains, we next examined the temporal profile of this change. Western blot analysis of brain lysates from WT and HDQ140/Q140 mice at age 1.5, 3, 5, and 10 months revealed that MAG levels in the striatum of HD mice began to decline at 3 months and progressively declined with age with age (**Figure 3A, 3B**). In the cerebral cortex, MAG reduction was first detected at 5 months (**Figure 3C, 3D**). In the striatum of HD mice, MAG loss preceded the reduction of MBP, MOBP, and MOG, which are localized at outer layers of myelin membranes. Levels of MOBP in HDQ140/Q140 mouse striata were transiently decreased at age 1.5 months but achieved to normal levels at age 3 months and then declined with age (**Figure 3A, 3B**). In the cerebral cortex, the levels of MAG, MOBP and MOG in HDQ140/Q140 mice were reduced at age 5 and 10 months, whereas MBP levels remained unaltered across all ages examined (**Figure 3C, 3D**). The level of NG2, a marker for OPCs (*43*), was normal in HDQ140/Q140 mice at all ages examined, but the level of Olig2, a transcription factor identifying the oligodendroglia lineage (*44*), was reduced in the striatum at 10 months (**Figure 3A – 3D**). The finding of normal NG2 levels in HD mouse brains suggests that the capacity of generating new oligodendrocytes is normally reserved in HD mouse brains. Consistent with immunohistochemistry studies (**Figure 2**), Western blot analysis revealed no difference in pNfH levels between WT and HDQ140/Q140 mice at all ages examined (**Figure 3A, 3C**), excluding axonal degeneration as a primary driver of loss of MAG and other myelin-related proteins. Finally, in heterozygous zQ175 mice, the reduction of MAG also preceded the loss of MBP, MOBP and MOG (**Figure 4A, 4B**). Combined, these results suggest that the loss of MAG protein in the striatum is an early phenotype of HD.

**Figure 3.**
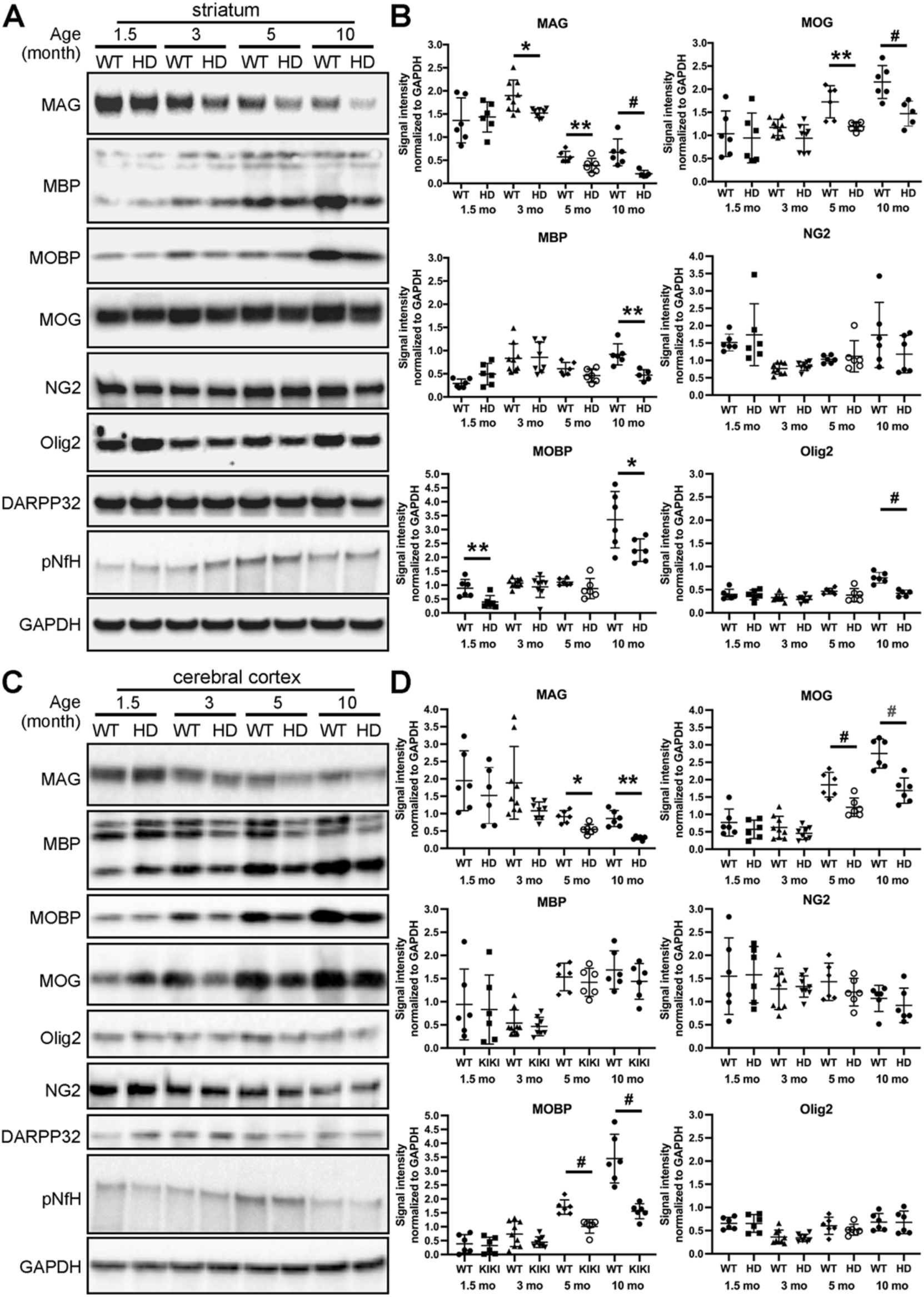
The reduction of MAG in the striatum is an early phenotype in HDQ140/Q140 mice and progressive with age. WT and HDQ140/Q140 mice were sacrificed at age 1.5, 3, 5, and 10 months. The brain was harvested from each animal and subjected to dissecting the striatum and the cerebral cortex. Western blot analysis of protein lysates prepared from striatal (**A**) and cerebral cortical (**C**) tissues with indicated antibodies followed by densitometrical analysis of the examined proteins in striatal (**B**) and in cerebral cortical (**D**) lysates. Photographs in (**A** and **C**) show blot analyses from one cohort of WT and HD mice at each age. Each symbol in (**B** and **D**) represents one animal. N=6 mice per genotype. Data are mean±SD. Statistical significance was determined by two-tailed unpaired Student’s t-test: * P<0.05; ** P<0.01; # P<0.001.

**Figure 4.**
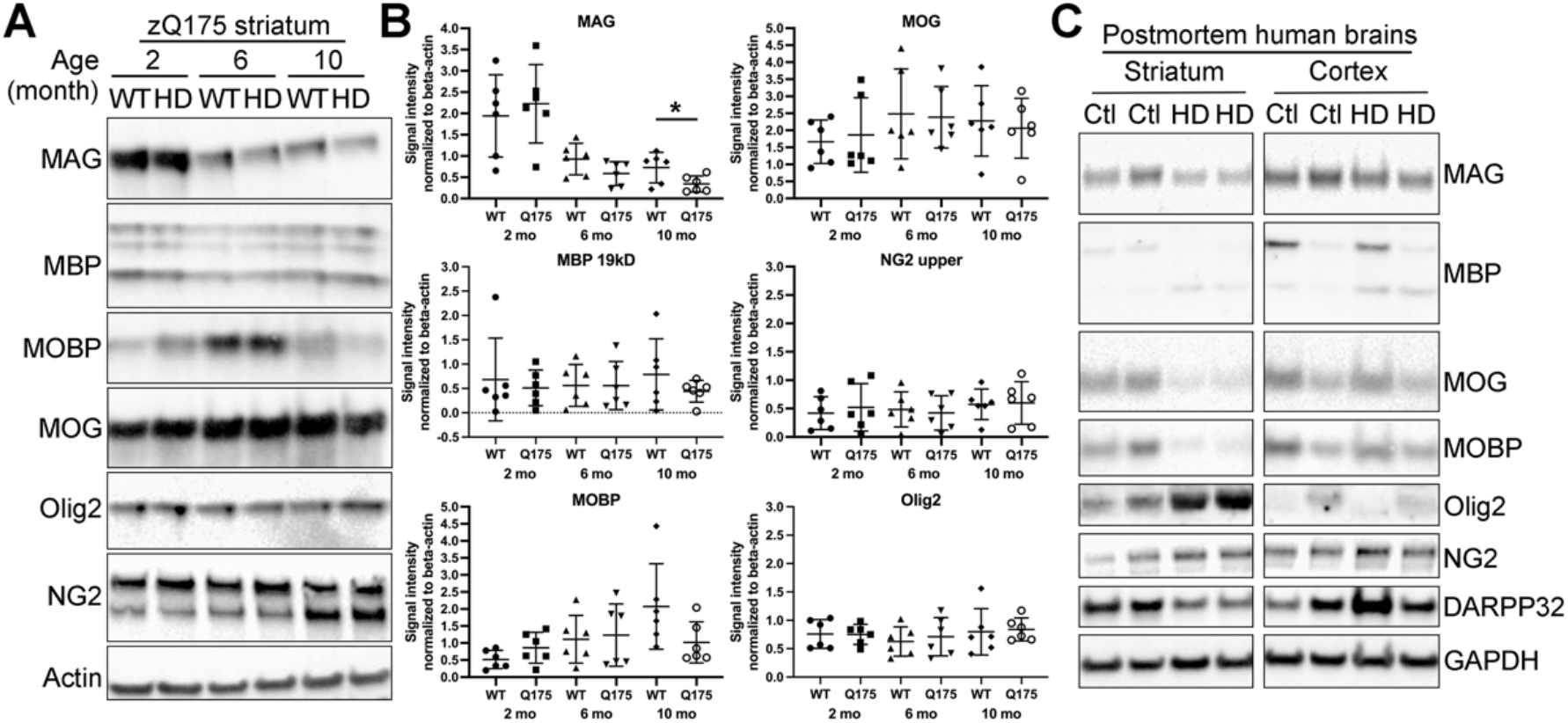
The reduction of MAG also occurs in heterozygous zQ175 mice and postmortem human HD brains. Western blot analysis of striatal lysates of zQ175 mice with indicated antibodies (**A**) followed by quantitative analysis of signal intensities of the examined proteins (**B**). Western blot analysis of striatal and cerebral cortical lysates of two patients with HD and two non-related healthy controls. Photographs in (**A**) show blot analyses from one cohort of WT and zQ175 mice at each age. Each symbol in (**B**) represents one animal. N=6 mice per genotype. Data are mean±SD. Statistical significance was determined by two-tailed unpaired Student’s t-test: * P<0.05.

### MAG reduction also occurs in human HD caudate

To determine whether MAG levels were reduced in the brain of patients suffering from HD, we evaluated postmortem human brain samples from two HD patients. Western blot analysis showed that MAG levels were reduced in neostriatal lysates and unaltered in cortical lysates (**Figure 4C**). MOG and MOBP levels were also diminished in HD neostriatal lysates, and the magnitude of their reduction was even greater than that of MAG reduction (**Figure 4C**). We observed that control neostriatal samples contained mainly the high-molecular-weight form of MBP, whereas the low-molecular-weight form of MBP was predominant in HD neostriatal samples (**Figure 4C**). A major difference among MBP isoforms is the inclusion/exclusion of exon II, which can direct MBP to the nucleus to influence oligodendroglia proliferation and morphological changes (*45*). It is unclear whether such a switch of MBP from high to low molecular weight isoforms in HD brains reflects a change in oligodendroglia proliferation and morphological remodeling. Nonetheless, these data indicate a substantial loss of myelin in the brain of these two HD patients. Consistent with prior observations of an increased abundance of oligodendroglia in postmortem human HD brains (*46, 47*), Olig2 levels were elevated in neostriatal lysates of both HD patients (**Figure 4C**). However, NG2 levels appeared unaltered in HD neostriatum and cerebral cortex (**Figure 4C**). These data suggest more oligodendrocytes produced, likely reflecting a compensatory attempt for myelin regeneration or repair, while the reserve pool of OPCs remains unchanged in human HD brains.

### MAG-labeled oligodendrocytes are newly generated ones

While MAG itself is not essential for the initial myelination, its deficiency delays oligodendrocyte maturation (*48, 49*). In human and mouse HD brains, more OPCs are committed to differentiation, but the maturation of oligodendrocytes is impaired (*6, 13*), suggesting that the abundance of new oligodendrocytes is increased in HD brains. BCAS1 was recently identified as a marker for new oligodendrocytes in mouse and human brains (*50*). To determine whether the reduction of MAG observed in HD mouse brains was associated with an altered abundance of new oligodendrocytes, brain sections from WT and HDQ140/Q140 mice were co-stained with antibodies against BCAS1 and MAG, respectively. Microscopic analysis showed that oligodendrocytes containing intense MAG signals in the soma expressed BCAS1 (**Figure 5A**), indicating that they are newly generated oligodendrocytes. As the expression of BCAS1 commences after OPCs are committed to differentiation and gradually declines as the maturation process progresses (**Figure 5B**), the temporal expression pattern of BCAS1 relative to other markers of the oligodendroglia lineage, e.g. PLP and MAG, has been used as an index of oligodendrocyte maturation stages (*50*). We therefore analyzed the intensity of MAG and BCAS1 immunoreactive signals in labeled oligodendrocytes in caudate putamen areas of WT and HD mouse brain sections (**Figure 5A**). Based on the relative signal strength of BCAS1 and MAG immunoreactivities, the labeled oligodendrocytes can be divided into 5 populations: 1) BCAS1-positive and MAG-negative, 2) high levels of BCAS1 with low levels of MAG, 3) similar levels of BCAS1 and MAG, 4) low levels of BCAS1 with high levels of MAG, and 5) BCAS1-negative and MAG-positive. We quantified these 5 populations of oligodendrocytes in both WT and HD brain sections (**Figure 5C**). The BCAS1-positive but MAG-negative oligodendroglia in subventricular zones were excluded in our analysis as the brain structures into which they migrate for myelinating axons are still unclear. Under these conditions, the overall frequency of BCAS1-positive oligodendrocytes in the caudate putamen was not different between WT and HD mice. However, a greater population of oligodendrocytes were BCAS1-positive and MAG-negative in HD mouse brains, whereas the major population of oligodendrocytes in WT mice expressed both MAG and BCAS1 (**Figure 5D**). Moreover, a subset of BCAS1-negative and MAG-positive oligodendrocytes was detected in HD mouse brains but rarely seen in WT mouse brains (**Figure 5D**). These findings indicate that newly generated oligodendrocytes in HD mouse exhibit delayed acquisition of MAG and impaired trafficking of MAG from the soma to processes, consistent with a maturation defect of newly generated oligodendrocytes in HD brains.

**Figure 5.**
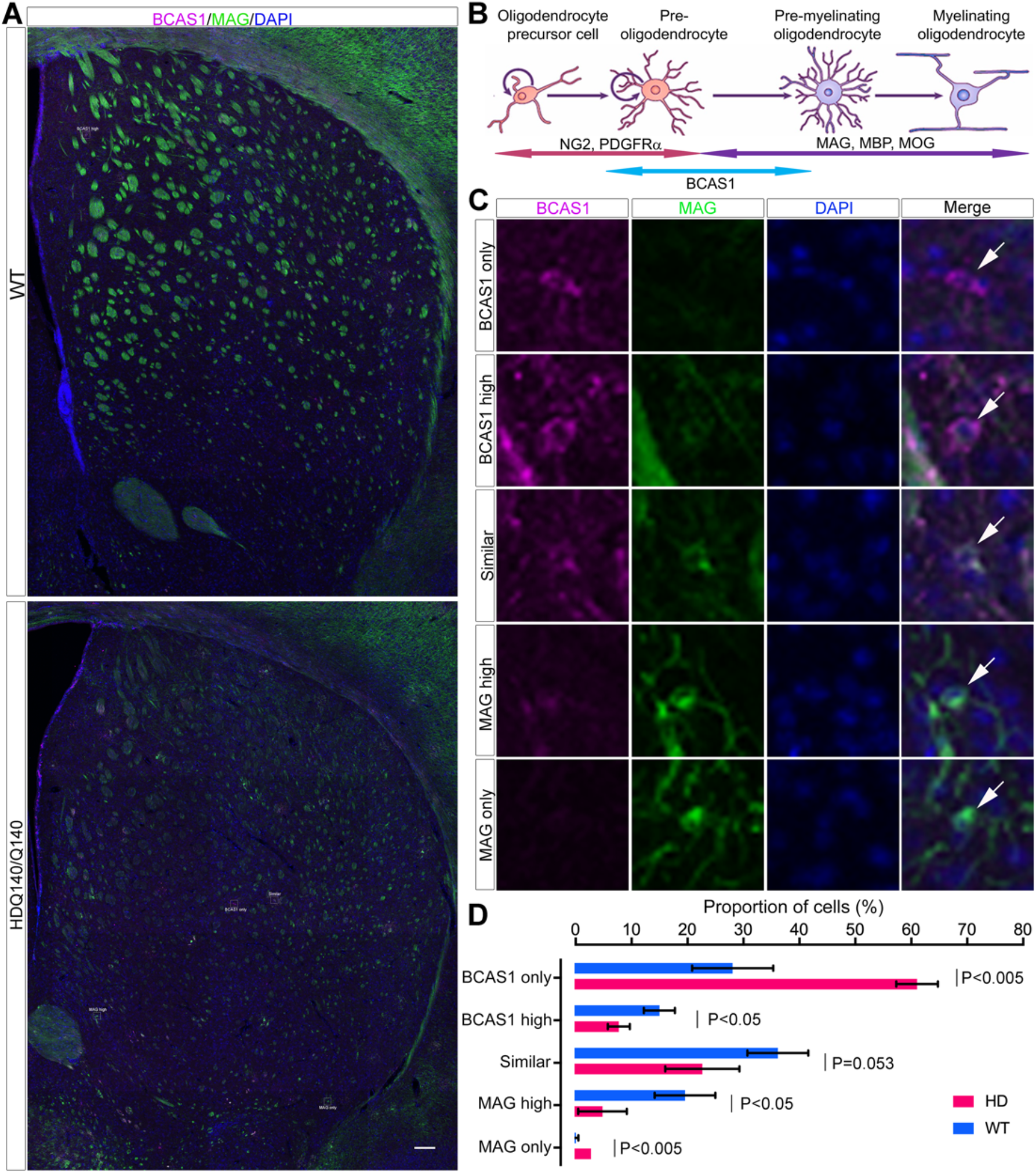
The oligodendrocytes showing MAG accumulation in the soma are newly generated ones. A series of 3 consecutive brain sections of each of WT and HDQ140/Q140 mice (n=3 mice per genotype) were processed for immunofluorescence labeling to detect MAG (green) and BCAS1 (violet). Cells in brain sections were identified by staining nuclei with DAPI. **A**) shown are confocal images of caudate putamen areas of a WT mouse brain section and an HD mouse brain section. Scale bar: 200 μm. **B**) Diagram of the temporal expression of marker proteins during the maturation of oligodendrocytes. Based on the temporal expression of BCAS1 relative to MAG, there should be 5 populations of oligodendrocytes in the brain. **C**) Shown are enlarged images of the corresponding boxed regions in images (**A**) to highlight the 5 populations of oligodendrocytes. **D)** Quantitative analysis of the 5 populations of oligodendrocytes. Data are mean±SD. Statistical significance was determined by two-tailed unpaired Student’s t-test (n=3 mice/genotype, 3 sections/mouse).

### MAG-accumulating structures originate from the late endosomal lysosomal compartment

To evaluate the subcellular distribution of MAG in BCAS1-positive oligodendrocytes, we captured and reconstructed z-stack confocal images from the caudate putamen of WT and HD mice (**Video S1**). MAG immunoreactivity appeared at punctate structures in perinuclear regions (**Fig 6A**). Quantitative analysis revealed that oligodendrocytes in HD mice exhibited a reduced frequency of small MAG-positive vesicles whereas an increased frequency of large MAG-positive perinuclear structures compared to oligodendrocytes in WT mouse brains (**Figure 6B**). This finding suggests that the generation of small vesicles transporting MAG away from large perinuclear structures was diminished in oligodendrocytes of HD mice and strengthens our idea that the targeting of MAG to myelin membranes was impaired in HD brains.

**Figure 6.**
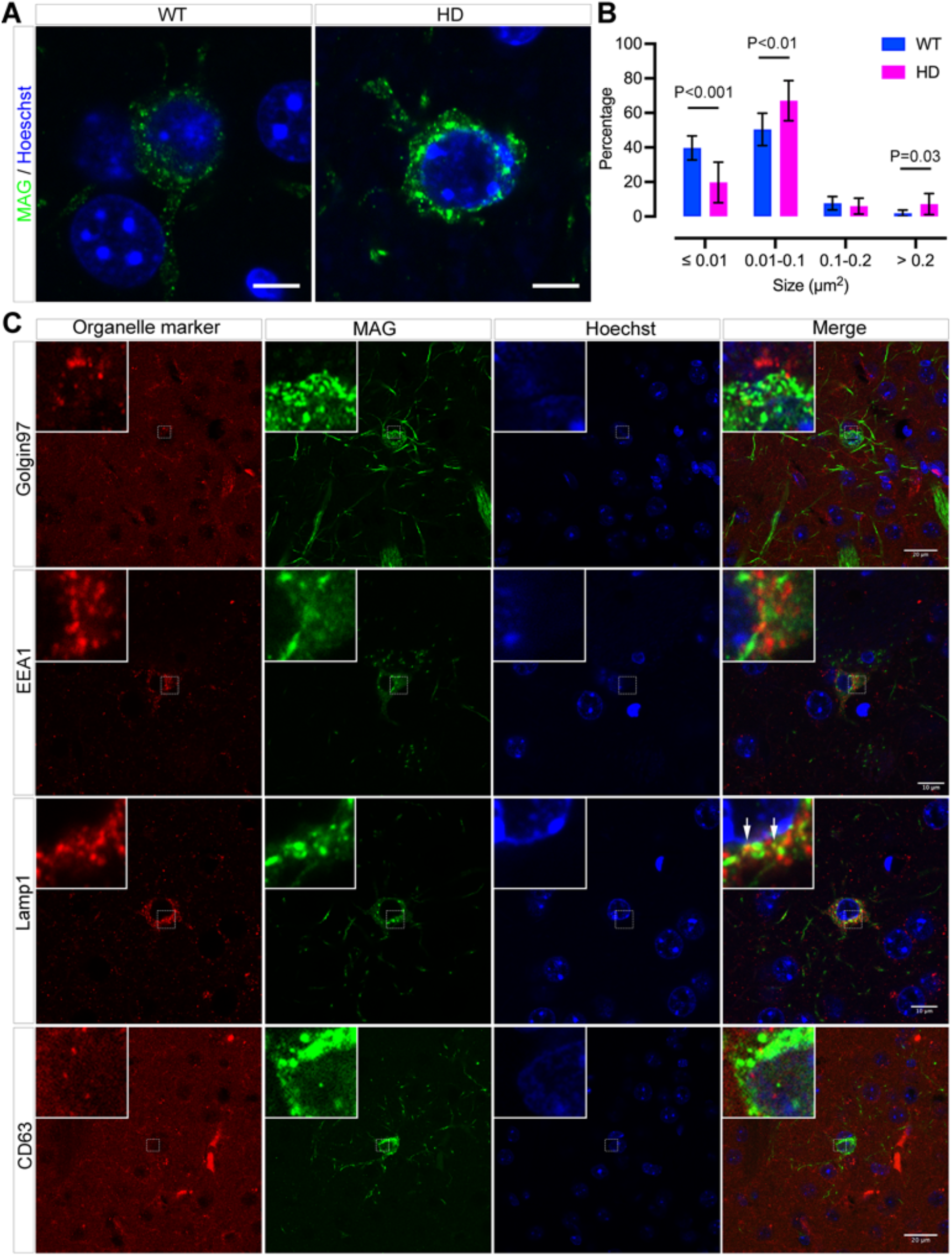
The MAG-accumulating structures in HD oligodendroglia are derived but distinct from late endosomes. Immunofluorescent labeling of WT and HDQ140/Q140 mouse brains to show MAG-containing structures in perinuclear regions. **A**) Shown are confocal images. Scale bars: 10μm. **B**) Quantitative analysis of the size of MAG-containing perinuclear structures in WT and HDQ140/Q140 mouse brains. Data are mean±SD. Statistical significance was determined by two-tailed unpaired Student’s t-test (n=3 mice/genotype, 3 sections/mouse). **C**) MAG-containing perinuclear structures are derived from late endosomal lysosomal compartments. A series of brain sections of HDQ140/Q140 mice were processed for labeling with antibodies for MAG combined with antibodies for Golgin97, EEA1, LAMP1, and CD63, respectively. Shown are confocal images. Arrows indicate structures labeled with both MAG and LAMP1.

We next sought to define the identity of MAG-accumulating perinuclear structures. Mutant Htt has been shown to cause an accumulation of vesicles at trans-Golgi networks and constrain the transport of early endosomes on microtubule cytoskeletons (*51, 52*). However, neither Golgin-97, which identifies trans-Golgi networks, nor early endosome antigen 1 (EEA1), which demarcates early endosomes, colocalized with MAG-containing perinuclear structures (**Figure 6C**), excluding trans-Golgi networks and early endosomes as the site of MAG accumulation. MAG is synthesized in the endoplasmic reticular and transported through the Golgi apparatus to the somatic surface, where it rapidly undergoes endocytosis and transiently passes through LAMP1-positive late endosomal lysosomal compartments before being targeted to myelin membranes (*53, 54*). In oligodendrocytes of neonatal rats, MAG has been detected in multivesicular bodies (*55*), from which intraluminal vesicles are secreted as exosomes (*56*). In various types of specialized cells, cell type-specific cargoes are stored in a specialized membrane compartment known as lysosome-related organelle, such as melanosomes in melanocytes (*57, 58*). To determine whether MAG-accumulating structures were exosomes and/or a lysosome-related organelle, brain sections were labeled with antibodies against MAG combined with antibodies against either CD63, a marker for exosomes, or LAMP1, a marker for late endosomes/lysosomes. Microscopic analysis showed that CD63 was absent in MAG-positive structures, but a small proportion of MAG-positive structures were labeled with LAMP1 (**Figure 6C**). Taken together, these data suggest that the MAG-accumulating structures are derived from the late endosomal lysosomal compartment.

## Discussion

White matter atrophy is a prominent pathological feature of HD, yet the underlying mechanisms are unclear. Studies in human and mouse HD brains suggest that the myelin undergoes pathologic changes prior to overt axonal damage, supporting a hypothesis of a de-myelination process that results in severe loss of the white matter in HD brains. In line with this idea, evidence of increased myelin repair attempts has been observed in HD brains. However, oligodendrocytes maturation and metabolism are impaired by the HD mutation, limiting the repair capacity. In this study, we identify MAG, a key adhesion and signaling molecule at the axon-myelin-interface, as an early and progressively affected protein in HD. In HDQ140/Q140 knock-in mice, MAG levels declined progressively in the caudate putamen first, then in the cortex, accompanied by reduced localization to fiber bundles and abnormal retention in the soma of newly generated oligodendrocytes. Although oligodendrocyte generation was preserved, their maturation was delayed: newly formed cells were slowed to acquire MAG expression and to traffic MAG from perinuclear compartments to myelin sheaths. Electron microscopy confirmed that MAG loss occurred at periaxonal myelin lamellae before axonal damage and overt myelin breakdown. The perinuclear structures enriched in MAG were enlarged in HD oligodendrocytes and did not contain protein markers for the Golgi apparatus (Golgin-97), early endosomes (EEA1), or exosomes (CD63). A minor portion were positive for LAMP1, suggesting derivation from the late endosome/lysosomal compartment.

How MAG is targeted to myelin-forming processes is not completely clear. MAG is a type-I transmembrane protein synthesized in the endoplasmic reticulum and transported through the Golgi apparatus to the plasma membrane of the soma, from where it undergoes endocytosis and transiently passes through LAMP1-positive late endosomal/lysosomal compartments *en route* to myelin-forming processes (*53, 54*). The altered vesicular distribution and size in HD, as well as the diminution at periaxonal membranes and fiber bundles suggests a bottleneck in this trafficking pathway.

The identity of the structures where MAG accumulates is unknown. Consistent with the finding that MAG transiently passes through LAMP1-positve late endosomal lysosomal compartments (*53-55*), a small portion of perinuclear MAG-positive structures contained LAMP1, suggesting that MAG-accumulating structures are derived but distinct from LAMP1-postive late endosomal lysosomal compartments. In myelinating oligodendrocytes of neonatal but not adult rats, MAG is detected at multivesicular bodies (*55*), which are discharged as exosomes. Yet, CD63, a marker used for identifying multivesicular bodies and exosomes (*56*), was not found on perinuclear MAG-labeled structures, indicating that they are not coalesced exosomes. Collectively, the perinuclear MAG-accumulating structures are likely a special organelle derived from LAMP1-positive late endosomal lysosomal compartments.

Lysosome-related organelles are a group of cell type-specific membrane compartments sharing features with endosomes and lysosomes. Most of lysosome-related organelles serve as a reservoir or storage compartments for cell type-specific cargoes and govern the secretion of such cell type-specific cargoes. Lysosome-related organelles have been found in a variety of cell types, e.g., melanosomes in pigment cells, Weibel-Palade bodies in endothelial cells, and dense granules in platelets (*57, 58*). By analogy, these MAG-positive structures may represent a previously undescribed oligodendrocyte-specific organelle for MAG storage and regulated release. Future studies are needed to identify this organelle.

The functional consequences of MAG loss in HD are likely multifaceted. Genetic inactivation of MAG does not affect the initial myelination but leads to increased unmyelinated axons in adult mice as well as decreased level of some myelin-related proteins (*19, 20, 59*). There is also evidence supporting a role for MAG in the differentiation of oligodendrocytes for re-myelination. MAG acts as a receptor mediating trophic signals to promote the differentiation of oligodendrocytes (*49*), and the maturation of oligodendrocytes is arrested when MAG trafficking out of LAMP1-positive late endosomal lysosomal compartments is compromised (*54*). Furthermore, the abrogation of MAG slows oligodendrocyte differentiation (*48*). These lines of evidence support the idea that the progressive loss and deficient targeting of MAG may contribute to both de-myelination and impaired maturation of oligodendrocytes for re-myelination in HD brains.

Oligodendrocytes are highly specialized cells. Their maturation requires the organization of the plasma membrane into the soma, process, and membrane sheet. The generation and maintenance of such membrane asymmetry relies on the vesicle machinery exploited for establishing the apical and basolateral domains in epithelial cells (*10, 11, 32, 34*). Mutant Htt disturbs vesicular trafficking in both secretory and endocytic pathways and compromises polarity formation (*37, 51, 52, 60*). As mHtt is expressed and formed aggregates in oligodendrocytes, vesicle trafficking defects observed in neurons and other HD cell types may also extend to oligodendrocytes. Consequently, altered MAG distribution may reflect a more general vesicle trafficking defect affecting multiple myelin-related proteins. In postmortem human and mouse HD brains, impaired oligodendrocyte maturation is accompanied with a global repression of oligodendroglia lineage genes and particularly the dysfunction of myelin regulatory factor (MYRF), a transcription regulator that drives the expression of myelin-associated proteins including MAG (*61*). In this regard, the delayed expression of MAG in HD oligodendrocytes may be partly due to impaired functions of MYRF and other gene transcription regulators.

## Conclusions

Our study demonstrates that HD mice present with an early and progressive loss of MAG. MAG loss occurs before overt myelin breakdown, axonal degeneration, and the reduction of proteins localized to the outer layers of the myelin membrane. In HD mouse brains, new oligodendrocytes were normally generated but showed delayed acquisition of MAG expression and impaired targeting of MAG from somatic structures to myelin processes. MAG retention in lysosome-related organelle points to a trafficking bottleneck in HD oligodendrocytes. Our study suggests that defective MAG targeting is a critical contributor to white matter pathology in HD.

## Supporting information

Un-cropped blots

## Data availability

All data are included in the article and supplementary information files.

## Acknowledgements

Cryo-sectioning and immunogold labeling were conducted with the aid of Diane Capen in the Microscopy Core of the Program in Membrane Biology, Massachusetts General Hospital.

## Author contributions

X.L. conceptualized the study, designed experiments, conducted data analysis, prepared figures, and wrote the original manuscript. A.B., E.S., and K.S. conducted experiments and data analysis, prepared figures, and revised the manuscript. Y.Y.L. analyzed immunohistochemistry and electron microscopic images and prepared the graphic diagram. T.P. and G.SV. provided research materials and revised the manuscript. M.D., K.KG. and N.A. revised the manuscript.

## Ethics approval and consent to participate

Not applicable.

## Consent for publication

Not applicable.

## Graphical representation

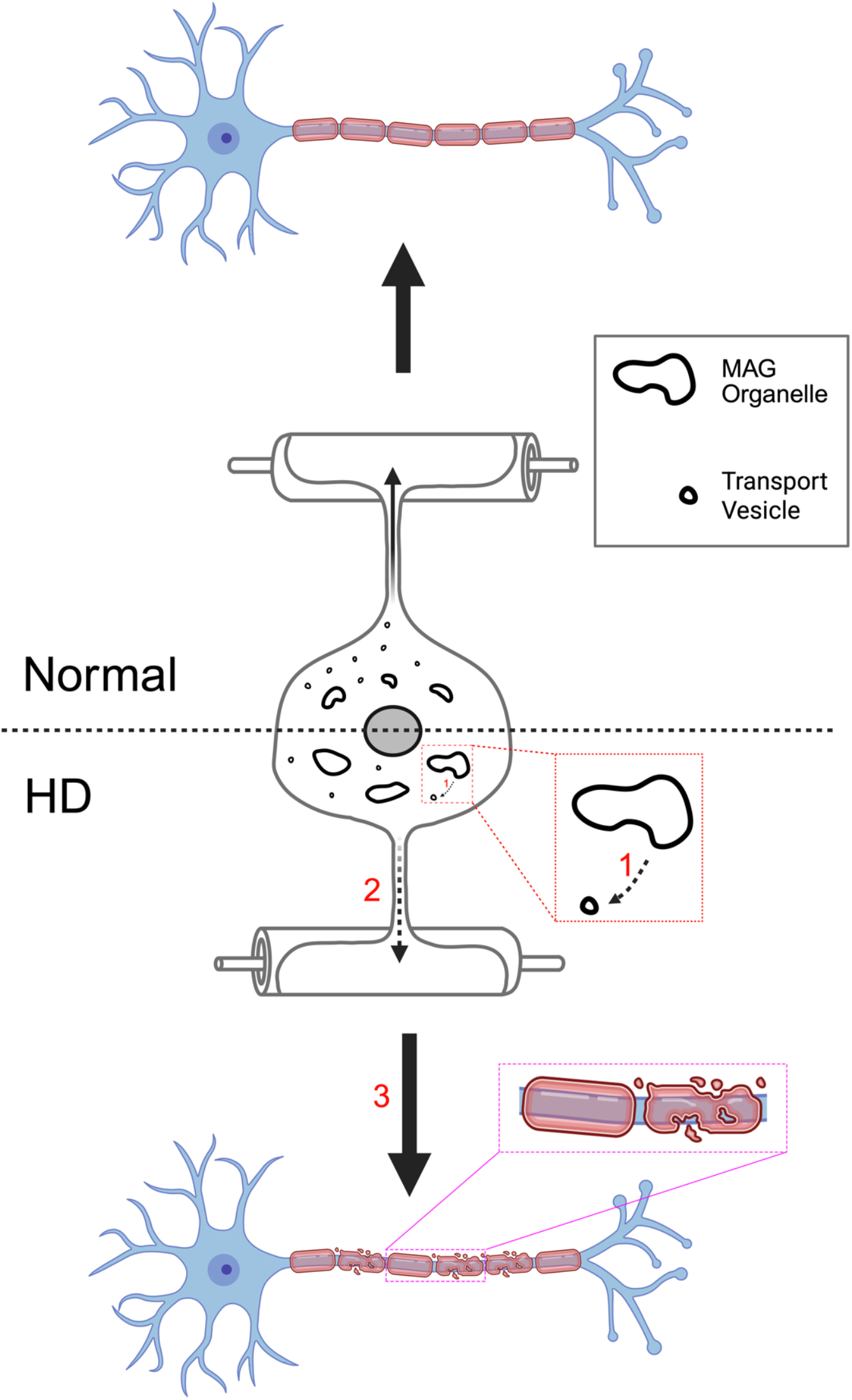

## Graphical representation

The expression of mutant huntingtin in oligodendrocytes impairs the generation of MAG-transporting vesicles from the MAG perinuclear compartment (Step-1), thereby reducing the delivery of MAG to myelin-forming processes (Step-2). MAG is essential for maintaining the periaxonal space; its decrease at myelin membranes causes the separation of the myelin sheath from the enwrapped axon and leads to de-myelination (Step-3). Additionally, the deficiency of MAG has been shown to significantly slow the differentiation of oligodendrocytes; a defect in its targeting to oligodendroglia processes may compromise the maturation of oligodendrocytes for repairing damaged myelin (Step-3). The graph was drawn with the bioRender software (www.biorender.com)

