## Supplementary material for "Early reduction and impaired targeting of myelin-associated glycoprotein to myelin membranes in Huntington’s disease": Un-cropped blots

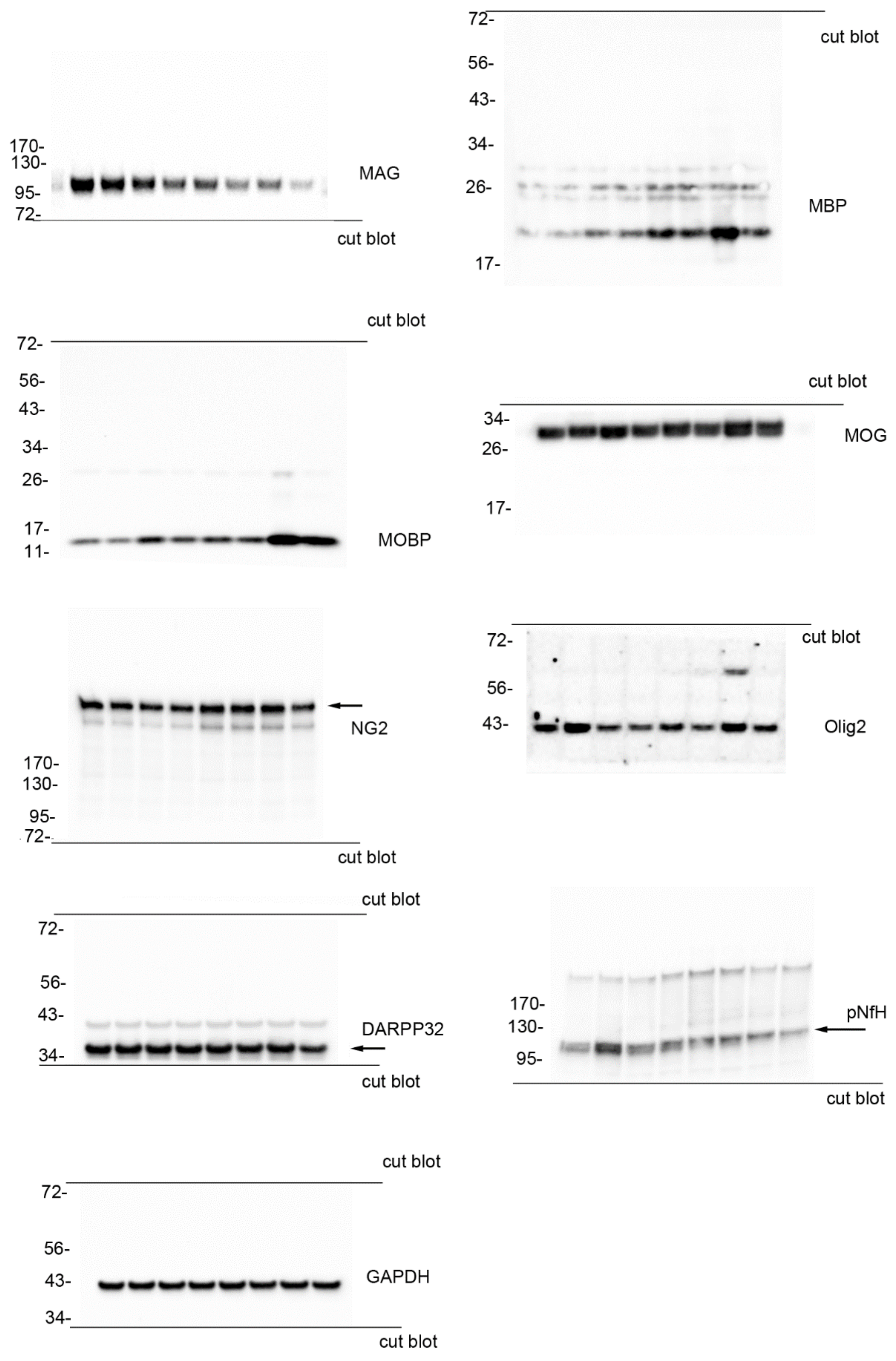


Full blots for **Figure 3A**. Blots were cut into horizontal strips as indicated to maximize the number of antibodies that could be analyzed per gel. DARPP32 blot was a reprobe of the GAPDH blot and residual signal is present.

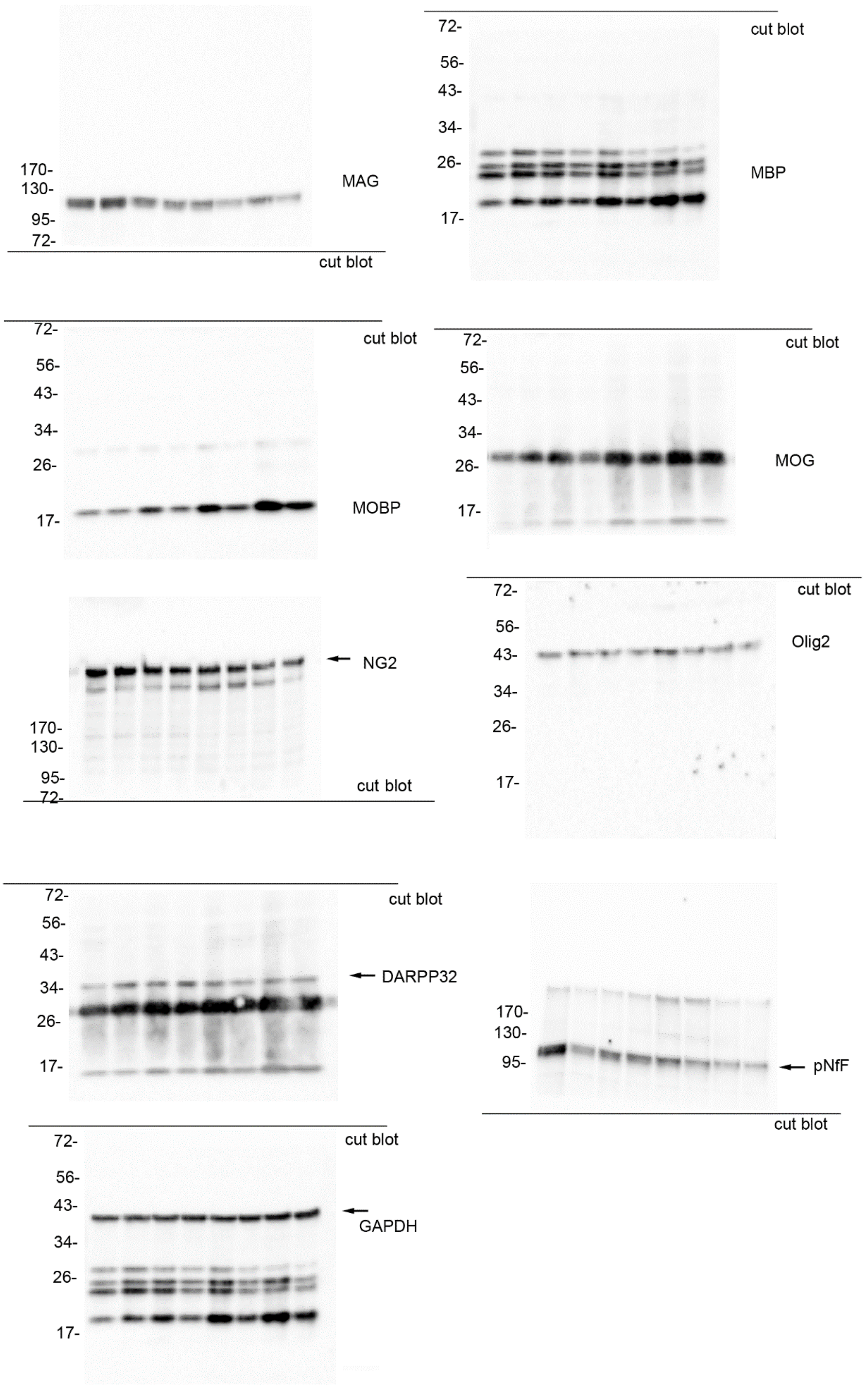


Full blots for **Figure 3C**. Blots were cut into horizontal strips as indicated to maximize the number of antibodies that could be analyzed per gel. DARPP32 blot was a reprobe of the MOG blot and GAPDH blot was a reprobe of the MBP blot and residual signals are present.

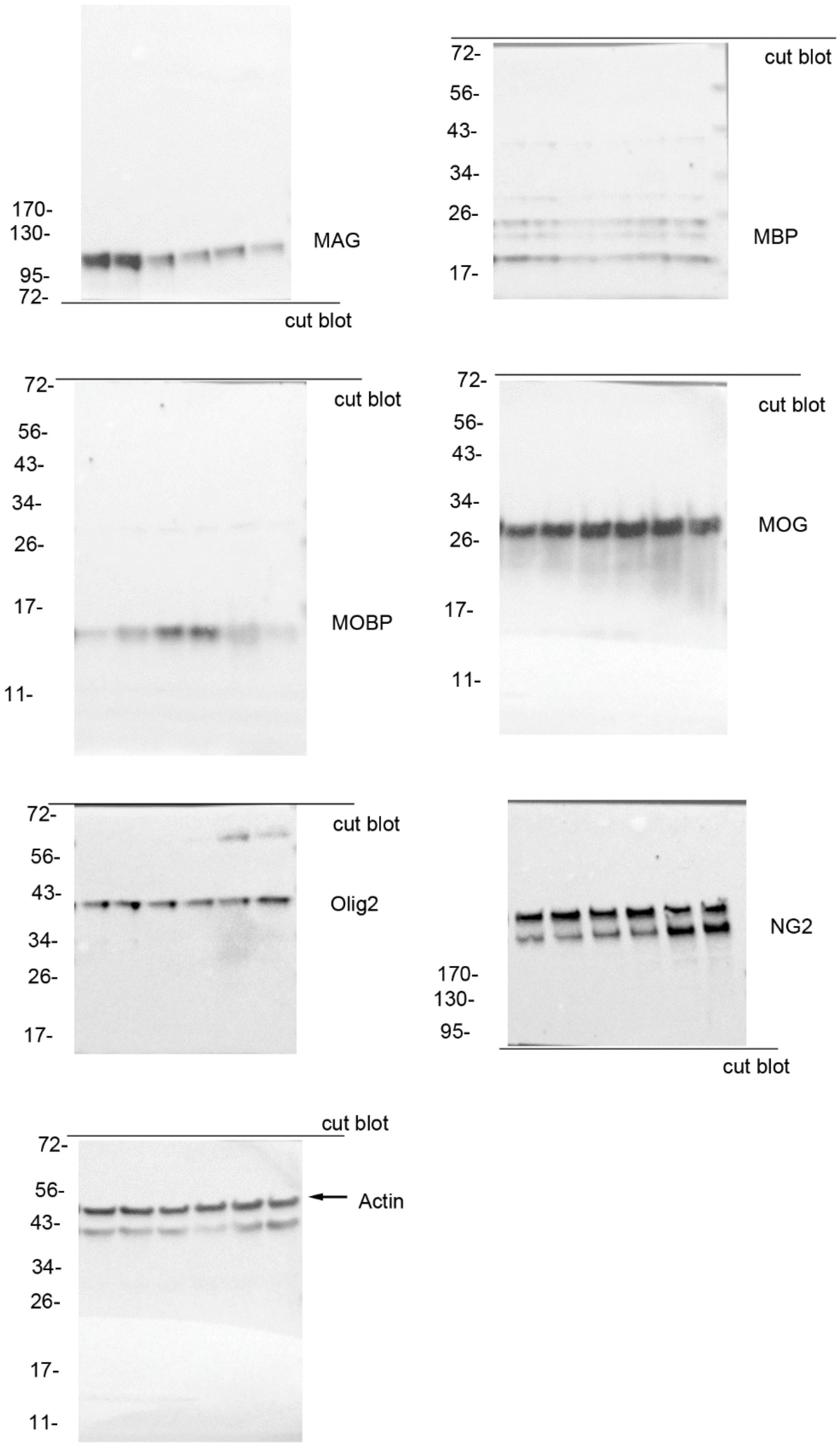


Full blots for **Figure 4A**. Blots were cut into horizontal strips as indicated to maximize the number of antibodies that could be analyzed per gel.

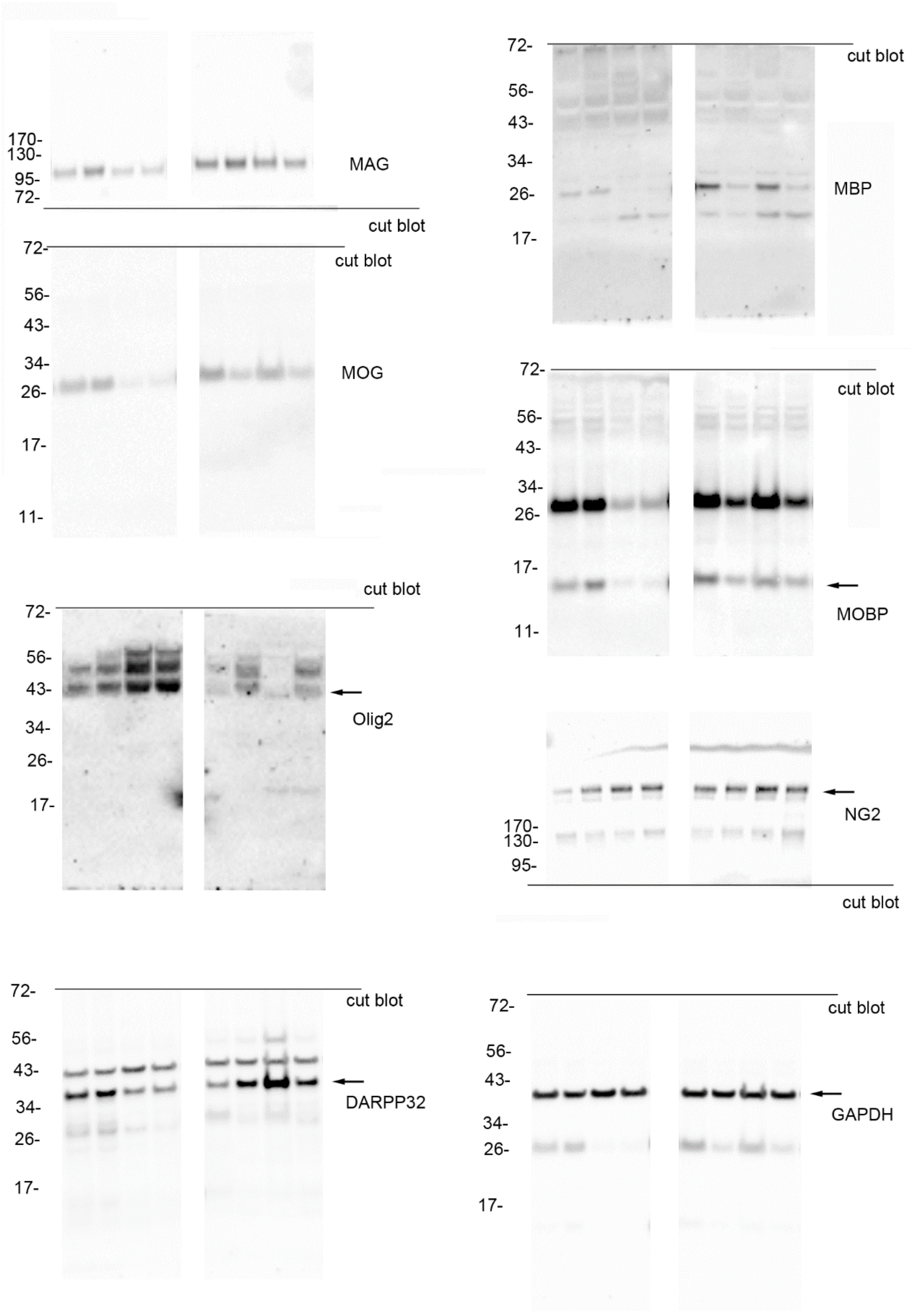


Full blots for **Figure 4C**. Blots were cut into horizontal strips as indicated to maximize the number of antibodies that could be analyzed per gel. MOBP and GAPDH blots were reprobes of the MOG blot and DARPP32 blot was a reprobe of GAPDH blot and residual signals are present.
