## Supplemental figure S1 and S2 for "Early reduction and impaired targeting of myelin-associated glycoprotein to myelin membranes in Huntington’s disease"

### Slide 1
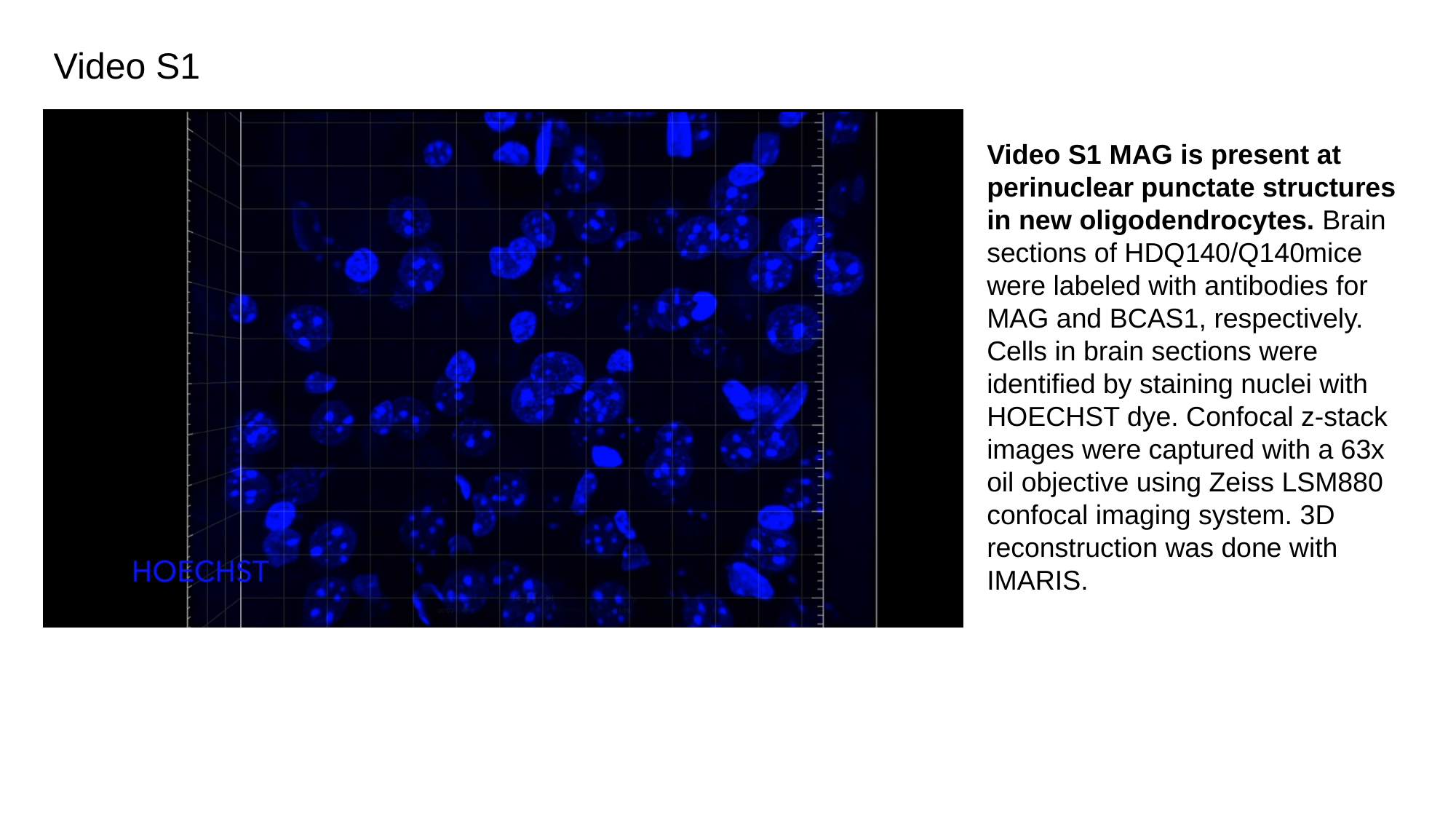

Video S1
Video S1 MAG is present at perinuclear punctate structures in new oligodendrocytes. Brain sections of HDQ140/Q140mice were labeled with antibodies for MAG and BCAS1, respectively. Cells in brain sections were identified by staining nuclei with HOECHST dye. Confocal z-stack images were captured with a 63x oil objective using Zeiss LSM880 confocal imaging system. 3D reconstruction was done with IMARIS.

### Slide 2
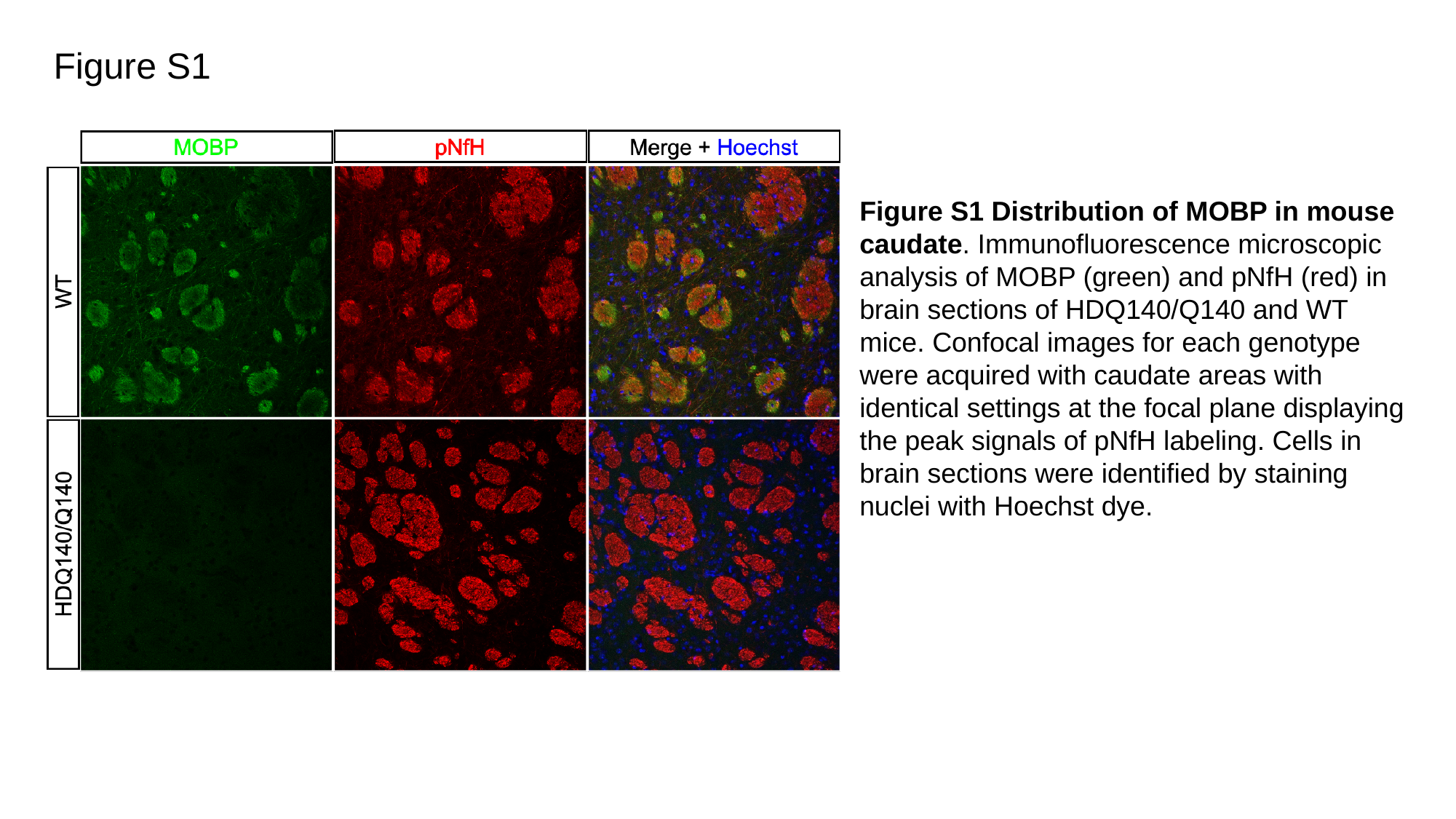

Figure S1
Figure S1 Distribution of MOBP in mouse caudate. Immunofluorescence microscopic analysis of MOBP (green) and pNfH (red) in brain sections of HDQ140/Q140 and WT mice. Confocal images for each genotype were acquired with caudate areas with identical settings at the focal plane displaying the peak signals of pNfH labeling. Cells in brain sections were identified by staining nuclei with Hoechst dye.
